# Effects of chronic intraperitoneal administration of the GABA_B_ receptor agonist 3-aminopropyl (methyl) phosphinic acid on food intake and body weight in rats

**DOI:** 10.64898/2026.01.18.700182

**Authors:** Ivor S Ebenezer

## Abstract

Previous research indicates that chronic intraperitoneal (ip) administration of the GABA_B_ receptor agonist baclofen reduces body weight gain in rats without altering daily food intake. The present study was undertaken to extend these observations by investigating the effects of chronic ip administration of the potent GABA_B_ receptor agonist 3-aminopropyl (methyl) phosphinic acid (SKF-97541) on daily changes on body weight and food intake in free feeding rats. The animals were injected ip once daily with SKF-97541 (0.1 mg / kg for 5 days, followed by 0.2 mg / kg for 8 days; Experiment 1) or SKF-97542 (0.4 mg / kg) for 17 days (Experiment 2). Control animals received physiological saline in both experiments. While daily food intake did not differ significantly between groups, the SKF-97541 group exhibited significant reductions in body weight gain compared to controls. These results extend previous findings and show that systemic administration of SKF-97541 suppresses weight gain independently of caloric intake, and lend further support to the hypothesis that GABA_B_ receptor agonists decrease body weight primarily by increasing metabolic rate.

## 1. Introduction

The role of GABA_B_ receptors in appetite stimulation is well-documented. Extensive research has established that both central (intracerebroventricular, icv) and systemic administration of the GABA_B_ receptor agonist baclofen increases food intake in free feeding and satiated animals, including pigs, rats, and mice (Ebenezer, 1990, Ebenezer and Baldwin, 1990, Ebenezer and Pringle, 1992, Ebenezer, 1995; Ebenezer and Patel, 2004; Higgs and Barber, 2004; Buda-Levin et al., 2005; Ebenezer and Prabhaker, 2007, Patel and Ebenezer, 2008a). These hyperphagic effects are mediated by central GABA_B._ receptors, which play a critical physiological role in regulating feeding behaviour (Ebenezer and Patel, 2004; Patel and Ebenezer, 2004). Pharmacological and immunohistochemical studies have identified the medial raphe nucleus, nucleus accumbens, and the arcuate nucleus of the hypothalamus as key regions involved in these responses (Decavel and Van den Pol, 1990, Wirtshafter et al., 1993, Stratford and Kelley, 1997, Ward et al., 2000, Backberg et al., 2003).

While most studies have focussed on acute administration and short-term feeding (Ebenezer and Pringle, 1992, Ebenezer, 1995, Higgs and Barber, 2004, Buda-Levin et al., 2005, Ebenezer and Prabhaker, 2007), investigations into chronic daily injections of baclofen (1 and 4 mg/kg, ip) in free feeding rats have revealed a complex regulatory profile. Specifically, while chronic baclofen treatment consistently elicited short-term (30 min) hyperphagia without the development of tolerance over 27 days, 24-hour food intake remained unchanged, suggesting accurate daily caloric regulation. However, the 4 mg / kg dose significantly reduced body weight, implying that activation of GABA_B_ receptors may influence food intake and body weight through distinct but different mechanisms. Similar findings with baclofen have been reported in the mouse (Ebenezer, 2005s). 3-Aminopropyl (methyl) phosphinic acid (SKF-97541) is a potent GABA_B_ receptor agonist (Seabrook et al., 1990; Froestl et al., 1995; Lehmann et al., 2011) that has recently been shown to stimulate short-term food intake in free-feeding rats (Ebenezer, 2026). However, its long term effects on food intake and body weight still need to be determined. The current study was therefore undertaken to investigate the effects of chronic administration of SKF-97541 on body weight and 24-hour food consumption in free feeding rats with the aim of extending and validating previous findings with baclofen (Ebenezer, 2025a; Ebenezer & Patel, 2010).

## 2. Methods and Material

The protocols used in this study were approved by the Ethical Review Committee at the University of Portsmouth, U.K. and carried out under licence granted by the United Kingdom Home Office.

### 2.1. Effects of repeated administration of SK-97541 on long-term (24 h) food intake and body weight gain in rats

#### 2.1.1 Experiment 1

Adult male Wistar rats (n=15; b. starting body weights: 255 – 335g; age = 9 weeks at start of experiment) were divided into 2 groups of similar body weights and housed in groups of 3 or 4 where they had free access to food (Food composition: (a) Percentage mass: protein 20%, oil 4.5%, carbohydrate 60%, fibre 5%, ash, 7% + traces of vitamins and metals, (b) Percentage energy: protein 27.3%, oil 11.48% and carbohydrate 61.2%, (c) Energy density: 3.600 kcal/g) and water.

The animals were kept on a 12h light-dark cycle (lights were turned on at 8.30 h and were turned off at 20.30 h). They given 2 training sessions when they were allowed free access to their normal pelleted food and water in experimental cages measuring 32 x 25 x 10 cm over a 24 h period. The food was presented to the rats in shallow cylindrical cups, as described previously (Ebenezer, 1990). During experimental sessions that followed, rats in Group 1 (n = 7) received saline and those in Group 2 (n = 8) received SKF-97541 (0.1 mg / kg for 5 days, followed by 0.2 mg / kg for the next 8 days). Saline and SKF-97541 were administered ip once daily at 14.00h and body weight measured prior to injections. The first injections was given to the rats on Day 0 of the experiment after measurement of body weights. The rats were returned to their home cages after injections except on experimental days 8 and 12 when the rats were placed separately in the experimental cages immediately after ip injection of saline (Group 1) or SKF-97541 (Group 2) with free access to food and water and cumulative food-intake measured 24 h later.

#### 2.12. Experiment 2

Adult male Wistar rats (n =12, starting body weight 390 - 440g) were treated in a similar manner to the rats in Experiment 1. During experimental sessions that followed, rats in Group 1 (n = 6) were injected ip with physiological saline and those in Group 2 (n = 6) injected ip with SKF-97541 (0.4 mg / kg) once daily at 14.00h and body weight measured prior to injections. The rats were returned to their home cages after injection except on experimental days 1, 8 and 16 when they were placed separately in the experimental cages immediately after ip injection of saline (Group 1) or SKF-97541 (Group 2) with free access to food and water and cumulative food-intake measured 24 h later.

### 2.2. Drugs

3-Aminopropyl (methyl) phosphinic acid (SKF-97541) was purchased from Sigma Biochemicals, Dorset, UK. The drug was dissolved in physiological saline solution (0.9% w/v, NaCl) to give an injection volume of 0.1 ml/100 g body weight. Physiological saline solution was used as the control.

### 2.3. Body Weight

The body weight data obtained for each rat were expressed as a percentage of the animal’s body weight recorded on Day 0 before receiving the first injection.

### 2.4. Statistics

The data from these experiments were analysed by two way mixed analysis of variance (ANOVA) with repeated measures on treatment and time (days) and by *post-hoc* t-test (Winer, 1971; see Discussion for further information on data analysis.

## 3. Results

### 3.1 Effects of repeated administration of SKF-97541 on long-term (24 h) food intake and body weight in rats

#### 3.11. Experiment 1

Even though two doses of SKF-97541 (0.1 mg / kg for 5 days, followed by 0.2 mg / kg for the next 8 days) were used in Experiment 1, the doses of the drug was not taken into account during the statistical analysis of the data (see Discussion for details).

The results on the effects of SKF-97541 on 24h cumulative food intake recorded following injections on experimental days 8 and 15 are illustrated in Fig. 1. Statistical analysis of the data revealed that there was no significant effects of treatment (F_(1,13)_ = 1.725, ns), time (F_(1,13)_ = 2.01, ns) and treatment x time interaction (F_(1,13)_ = 1.98, ns). Thus, the 24 h cumulative food intake following administration of SKF-97541 on days 8 and 15 was not significantly different from those of control rats.

**Figure 1.**
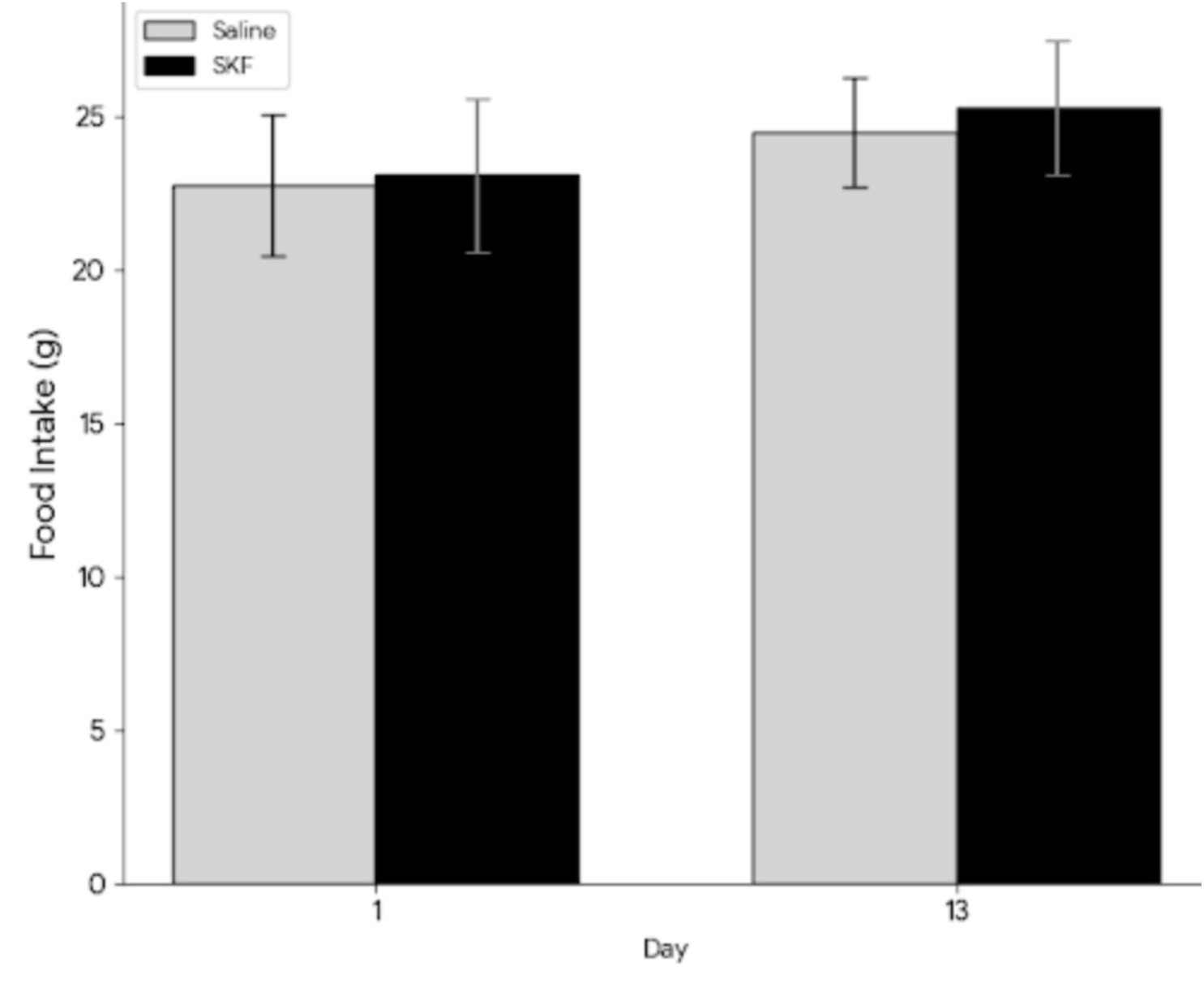
Effects of chronic intraperitoneal administration of physiological saline or SKF-97541 (0.1 mg / kg for 5 days followed by 0.2 mg / kg for the next 8 days) on 24 h food intake following injections on Days 8 and Day 13. See text for further details.

The saline treated control rats displayed a steady increase in body weight compared with their starting weights. However, the rats treated with SKF showed significant decreases in their body weights compared with saline-treated controls over the 14 day period of the experiment. (Fig. 2). Statistical analysis of the data revealed significant effects of treatment (F _(1,13)_ = 10.23 P<0.01), time (F _(13,169)_ = 3.24 P<0.01 and treatment x time interaction (F _(13,169)_ = 2..68 P<0.05). Interestingly, SKF-97541 decreased mean body weights of the rats to below their starting weights during the first 3 days (see Fig. 2).

**Figure 2.**
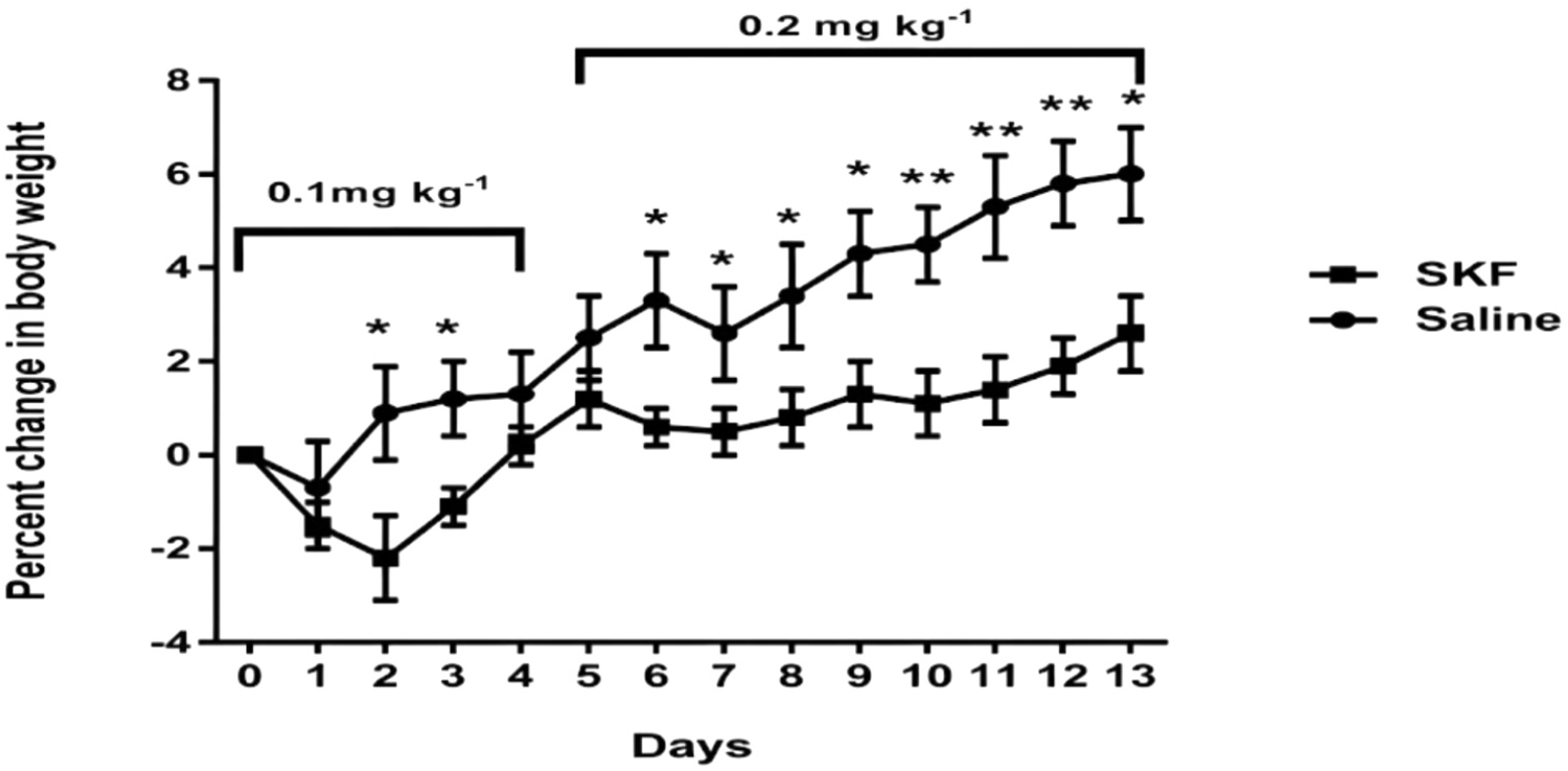
Effects of daily intraperitoneal injections of physiological saline or SKF-97541 (0.1 mg / kg for 5 days followed by 0.2 mg / kg for the next 8 days) on body weight in rats.. See text for further details.Vertical lines represent + or - S.E.M. *P<0.05 (SKF-97541 group vs saline control group).

The injections of SKF-97541 caused mild ataxia and sedation in the rats during the first 2 days. They were most apparent in the first 30 to 60 min. Tolerance developed to these effects with repeated administration of the drug.

#### 3.12. Experiment 2

The results on the effects of SKF-97541 (0.4 mg / kg) on 24h cumulative food intake recorded following injections on experimental days 1, 8 and 15 are illustrated in Fig. 3. ANOVA revealed that there was no significant effects of treatment (F_(1,13)_ = 1.725, ns), time (F_(2,34)_ = 2.01, ns) and treatment x time interaction (F_(2,34)_ = 1.98, ns) and indicate that the 24 h cumulative food intake following administration of SKF-97541 on days 1, 8 and 15 was not significantly different from those of control rats.

**Figure 3.**
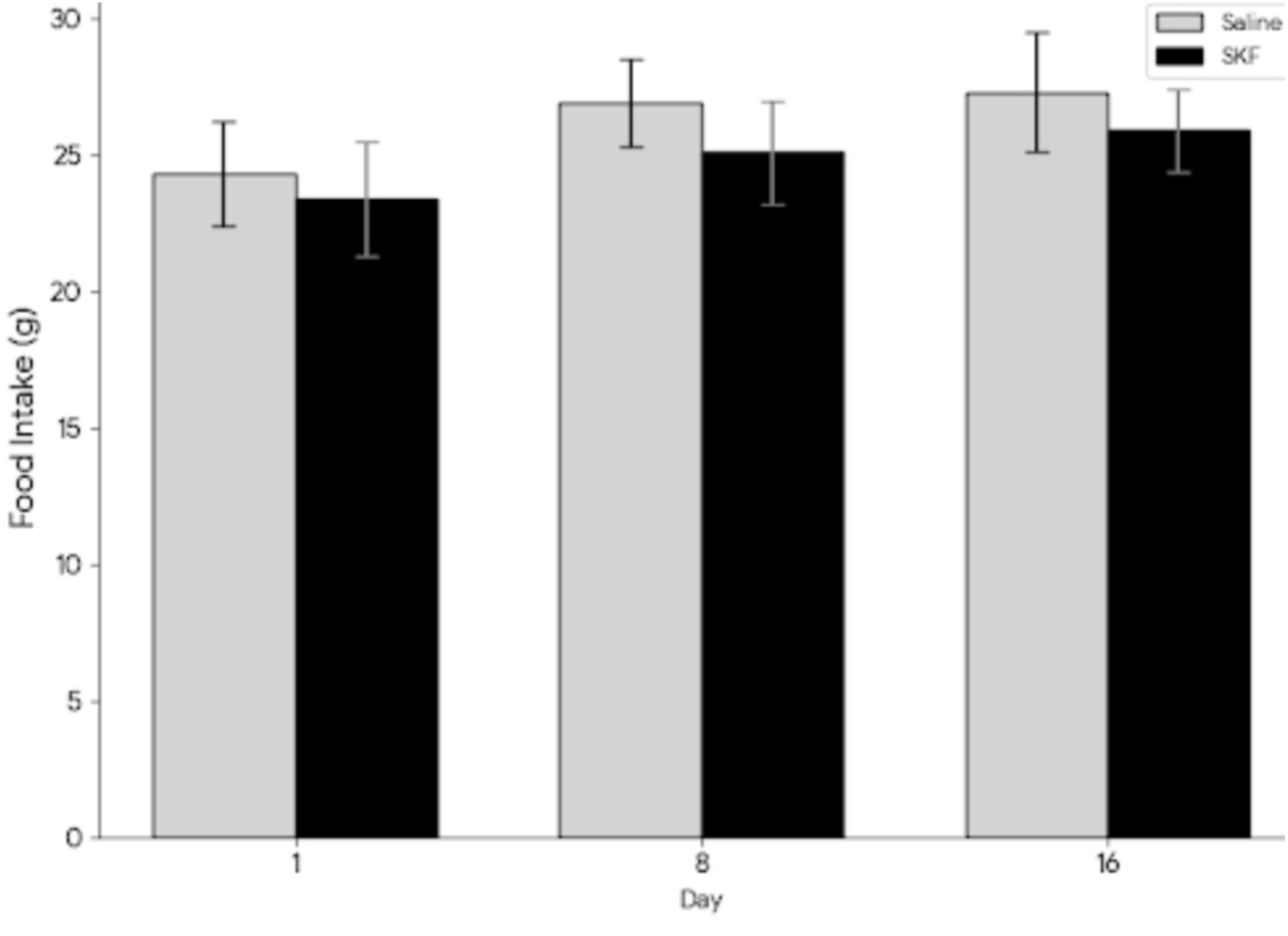
Effects chronic intraperitoneal administration of physiological saline or SKF-97541 (0.4 mg / kg for 17 days) on 24 h food intake following injections on Days 1, 8 and Day 16.. See text for further details.

The daily changes in body weight for the rats treated with physiological saline and SKF-97541 are shown in Fig. 4. ANOVA of the body weight data revealed that there were significant effects of treatment (F _(1,10)_ = 42.87, P<0.01), time (F _(16, 160)_ = 12.42 P<0.01 and treatment x time interaction (F _(16,160)_ = 28.68, P<0.01). The SKF-97541 (0.4 mg / kg) group exhibited significant reductions in body weight compared to the saline group (see Fig. 4). The saline treated control rats showed a steady increase in their daily body weights during the trial. However, the SKF-97541 treated animals displayed decreases in their daily body weights that were below their starting weights starting in day 1 and continuing until day 17 (see Fig. 4). By the end of the trials (day17), the rats in the saline group had gained an average of 3.08 % in body weight compared to their starting weight while the animals in the SKF-97541 group had lost an average of 3.87%.

**Figure 4.**
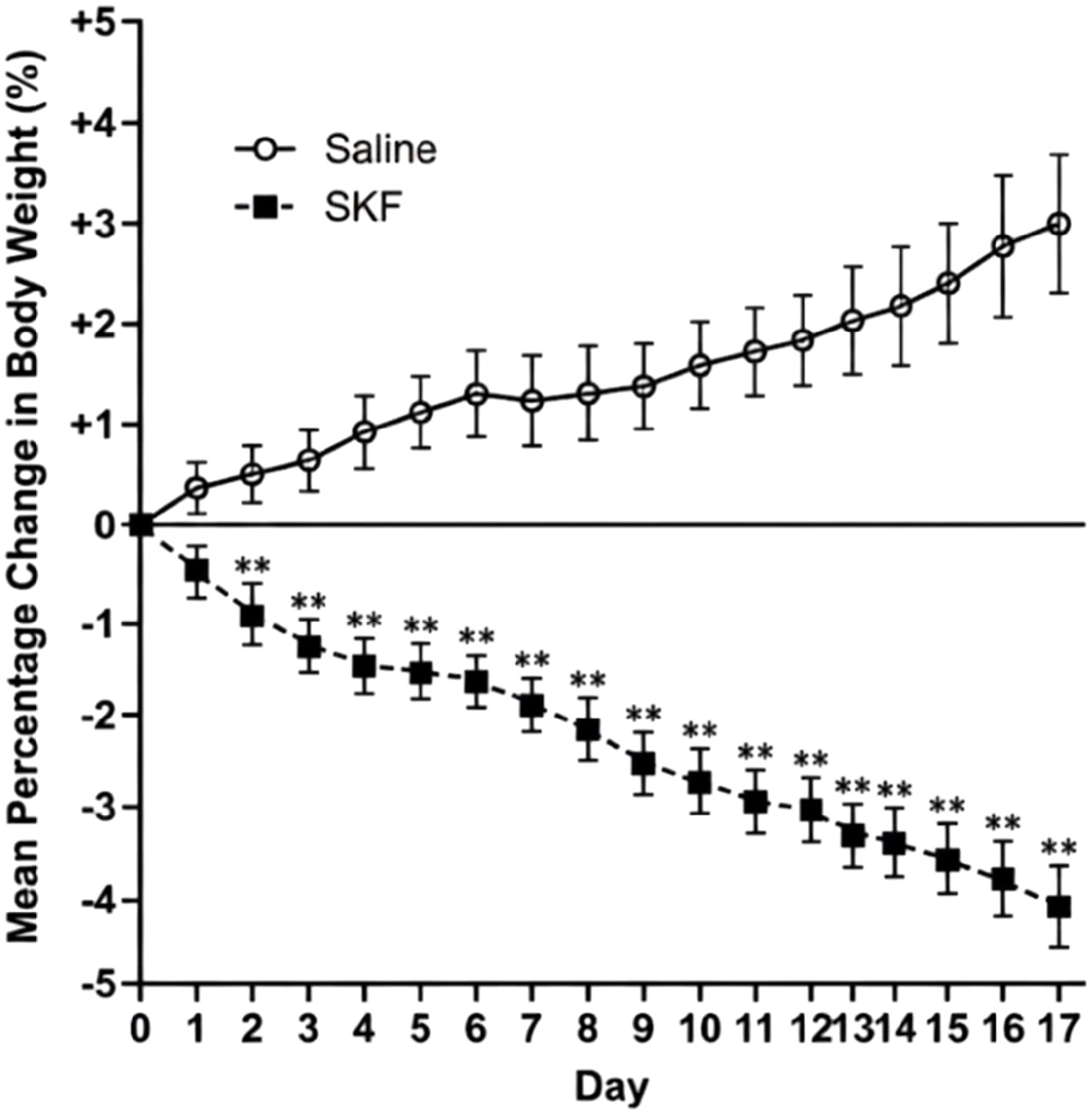
Effects of daily intraperitoneal injections of physiological saline or SKF-97541 (0.4mg / kg) over a period of 17 days on body weight in rats.. See text for further details.Vertical lines represent + or - S.E.M. **P<0.01 (SKF-97541 group vs saline control group).

The injections of SKF-97541 caused ataxia and sedation in the rats that were most apparent during the first 30 to 60 min. Tolerance developed to these effects with repeated administration of the drug.

## 4. Discussion

It has recently been reported that ip administration of the potent GABA_B_ receptor agonist SKF-97541 (3-aminopropyl (methyl) phosphinic acid) increases short-term food intake in free-feeding rats (Ebenezer, 2026). Building on this, the primary goal of the present study was to investigate whether chronic administration of SKF-97541 could reduce body weight in free-feeding rats without significantly altering long-term (24-hour) food intake, mirroring effects previously observed with the GABA_B_ agonist baclofen.

In the first experiment, rats received daily ip injections of SKF-97541, starting with a lower dose of 0.1 mg/kg for 5 days, followed by 0.2 mg/kg for an additional 8 days. This escalating dosing regimen was selected because acute administration of SKF-97541 produces dose-dependent behavioural side effects, primarily ataxia and sedation (Ebenezer, 2026). The initial lower dose helped minimise these effects and allowed the animals to acclimate to the drug, with rapid tolerance to the overt behavioural side effects developing upon repeated administration. Since the study aimed to assess the overall impact of chronic SKF-97541 treatment on 24-hour food intake and body weight, the data were analysed without separating the two dose phases. The results (Figs. 1 and 2) confirmed and extended previous findings with baclofen (Ebenezer, 2010), showing no significant differences in 24-hour food intake between SKF-97541-treated rats and saline controls. However, while saline-treated animals gained weight normally throughout the trial, those receiving daily SKF-97541 exhibited significant reductions in body weight gain. Notably, the mean body weights of the SKF-97541 group fell below their starting levels during the first three days (Fig. 2).

A second experiment was undertaken to investigated the effects of a higher daily dose of SKF-97541 (0.4 mg/kg) administered over 17 days. Previous work had shown that this dose does not significantly affect short-term (120-min) food consumption in free-feeding rats, largely due to pronounced behavioural side effects such as ataxia and sedation (Ebenezer, 2026). However, tolerance develops after repeated dosing over 5 days, after which the 0.4 mg/kg dose significantly increases short-term food intake (Ebenezer, 2026). In the present study, rats initially displayed similar side effects (mainly in the first 30–60 minutes post-injection), but tolerance developed after a few days. SKF-97541 had no significant effect on 24-hour food intake (measured on Days 1, 8, and 16; Fig. 3). Interestingly, the initial behavioural side effects on Day 1 did not impair overall 24-hour food consumption, suggesting the presence of compensatory mechanisms regulating daily intake (see Patel and Ebenezer, 2010).

Despite the lack of effect on food intake, saline-treated animals continued to gain weight normally, whereas those receiving daily SKF-97541 (0.4 mg/kg) showed significant reductions in body weight gain (Fig. 4). The divergence between groups was immediate and sustained, with the SKF-97541 group consistently falling below their starting weights throughout the experiment. By Day 17, saline-treated rats had gained an average of 3.08% in body weight relative to baseline, while the SKF-97541 group had lost an average of 3.87%.

It is noteworthy that rats treated with the higher dose of SKF-97541 (0.4 mg / kg) continued to lose weight, falling below their initial body weights throughout the 17-day trial (Fig. 4), whereas animals receiving lower doses of SKF-97541 (0.1 mg / kg followed by 0.2 mg/kg) showed only initial weight loss during the first three days, followed by gradual weight gain that eventually exceeded their starting weights (see Fig. 2). Comparable effects on body weight were previously observed with a 4 mg / kg dose of baclofen (Patel and Ebenezer, 2010, Ebenezer, 2025b). These differences between the 0.4 mg / kg dose of SKF-97541 and both the lower SKF-97541 doses and the 4 mg / kg baclofen dose likely stem from the relative pharmacological potency at GABA_B_ receptors. Current evidence indicates that SKF-97541 is at least ten times more potent than baclofen (Piqueras and Martinez, 2004; see also Seabrook et al., 1990). Accordingly, the present study used SKF-97541 at doses approximately tenfold lower than those of baclofen previously shown to increase food intake or reduce body weight gain during chronic administration (Ebenezer and Pringle, 1992, Ebenezer, 1995, Patel and Ebenezer, 2008a,b, Ebenezer and Patel, 2010). The 0.4 mg / kg dose of SKF-97541 thus appears substantially more potent as a GABA_B_ receptor agonist than 4 mg/kg baclofen, as reflected by the greater weight loss during chronic treatment and the more severe initial behavioural side effects.

It is also worth noting that SKF-97541 exhibits weak antagonist activity at GABA_A-ρ_ receptors (formerly known as GABA_C_ receptors; Naffaa et al., 2017). However, since these receptors are primarily localised in the retina, this property is unlikely to have influenced feeding behaviour or body weight changes in the present study.

The mechanisms underlying the body weight reduction observed with repeated administration of the GABA_B_ receptor agonists SKF-97541 and baclofen in rodents remain unclear. Notably, chronic treatment consistently lowers body weight without substantially altering daily food intake, suggesting that the effect is mediated by increased metabolic rate or energy expenditure rather than reduced food consumption (Ebenezer, 2025a,b, Patel and Ebenezer, 2011).

An early hypothesis involving central activation of the sympathetic nervous system (SNS) leading to brown adipose tissue (BAT) thermogenesis was mooted as a possible mechanism by which chronic administration of baclofen increased metabolic rate to decrease body weight (Patel and Ebenezer, 2010). It was demonstrated that microinjections of baclofen into the ventromedial nucleus of the hypothalamus (VMH) in anaesthetised rats activated BAT. This effect was blocked by the β-adrenoceptor antagonist propranolol, suggesting SNS outflow from central pathways to BAT (Rothwell et al., 1985, Addae et al., 1986). However, more recent evidence challenges this SNS-BAT model as the primary mechanism involved. Chronic administration of baclofen produced sustained weight loss without long-term changes in food intake, yet pretreatment with propranolol failed to prevent or reduce the effect (Ebenezer, 2005b). This findings suggest that enhanced sympathetic nervous system activation of BAT is unlikely to be the prime mechanism involved. However, there are a number of other possible ways by which chronic administration of GABA_B_ receptor agonist may decrease body weight., as discussed below.

It has been demonstrated that GABA_B_ receptors are present on BAT cells and that GABA_B_ receptor agonists act directly on BAT adipocytes to induce thermogenesis. (see Ikegami et al., 2018). Thus, SKF-97541 and baclofen may stimulate local thermogenesis by enhancing uncoupling protein-1 (UCP-1) activity in mitochondria, dissipating energy as heat without requiring sympathetic innervation. Furthermore, repeated baclofen administration could induce “browning” of white fat, a process where white fat cells acquire characteristics of brown fat, enhancing their metabolic activity and calorie burning characteristics (Machado et al., 2022) . This “browning” can be triggered by various factors, including cold exposure, exercise, and certain nutrients, such as omega-3 fatty acids (see Machado et al., 2022). Interestingly, preliminary results from our laboratory indicate that repeated daily administration of baclofen to free feeding rats increases browning of white fat (Bains and Ebenezer, unpublished results). Therefore, the weight loss observed with SKF-97541 may result from a combination of direct effects on BAT and the induction of browning of white fat.

It is also possible that GABA_B_ receptor agonists act centrally to induced reductions in body weight. Immunohistochemical and pharmacological evidence indicates that GABA_B_ receptors are found on the orexigenic neuropeptide Y (NPY) and the anorexigenic proopiomelanocortin (POMC) neurones in the arcuate nucleus of the hypothalamus (see Backberg et al., 2003, Cansell et al., 2012, Jasi and Bruning, 2022). These neuronal populations play a central role in the regulation of energy homeostasis (Morton et al., 2006, Sato et al., 2007, Haung et al., 2021). Therefore, it is possible that SKF-97541 may modulate the activity of NPY and/or POMC neurones in the actuate nucleus, potentially contributing to alterations in energy balance and body weight reduction.

In conclusion, the findings of this study extends previous observations with baclofen (Patel and Ebenezer, 2010, Ebenezer, 2025a,b) to another, more potent, GABA_B_ receptor agonist and shows that chronic ip injections of SKF-97541 decreases body weight that is not related to suppression of food intake. The weight loss is most probably the result of a combination of hypothalamic modulation, direct thermogenic stimulation, and the browning of white adipose tissue. However, further research is necessary to elucidate the precise mechanisms underlying the effects of drugs such as baclofen and SKF-97541 on body weight. These findings further support the potential therapeutic use of GABA_B_ receptor agonists in the treatment of obesity. This suggestion is supported by a preliminary open-label clinical trial that indicates that baclofen can reduce body weight in obese human subjects (Arima and Oiso, 2010).

## Acknowledgements

I wish to thank Taiwo and Asif for assistance in recording body weight for Experiments 1.

## Notes

### Competing Interest Statement

The authors have declared no competing interest.

